# *F5N* : Nanopore Sequence Analysis Toolkit for Android Smartphones

**DOI:** 10.1101/2020.03.22.002030

**Authors:** Hiruna Samarakoon, Sanoj Punchihewa, Anjana Senanayake, Roshan Ragel, Hasindu Gamaarachchi

## Abstract

*F5N* is the first ever Android application for nanopore sequence analysis on a mobile phone, comprised of popular tools for read alignment (*Minimap2*), sequence data manipulation (*Samtools*) and methylation calling (*F5C/Nanopolish*). On NA12878 nanopore data, *F5N* can perform a complete methylation calling pipeline on a mobile phone in ∼15 minutes for a batch of 4000 nanopore reads (∼34 megabases). *F5N* is not only a toolkit but also a framework for integrating existing C/C++ based command line tools to run on Android. *F5N* will enable performing nanopore sequence analysis on-site when used with an ultra-portable nanopore sequencer (eg: MinION or the anticipated smidgION), consequently reducing the cost for special computers and high-speed Internet.

**Availability and implementation:** *F5N* Android application is available on Google Play store at https://play.google.com/store/apps/details?id=com.mobilegenomics.genopo&hl=en and the source code is available on Github at https://github.com/SanojPunchihewa/f5n.

## INTRODUCTION

Ultra-portable single-molecule real-time sequencers such as MinION introduced by Oxford Nanopore Technologies (ONT) measure the ionic current when a DNA strand passes through a biological nanopore [1]. The sequencer periodically outputs a group of reads in the form of raw current signals (packed into a .*fast5* file) which are subsequently base-called (to a .*fastq* file) on a laptop or an ultra-portable ONT’s MinIT. Therefore, sequencing and base-calling processes have become portable but sequence analyses such as sequence alignment and genome polishing are still not. Currently, the sequence analysis is either performed using a cloud service (that involves uploading data over high-bandwidth network) or using dedicated high-end server computers, both of which are not synonymous with ultra-portable sequencing. We present the first ever mobile nanopore DNA sequence analysis toolkit *F5N*, which compacts popular DNA analysis tools to an Android mobile phone application to make sequence analysis fully portable (Fig. 1A).

**Fig. 1.**
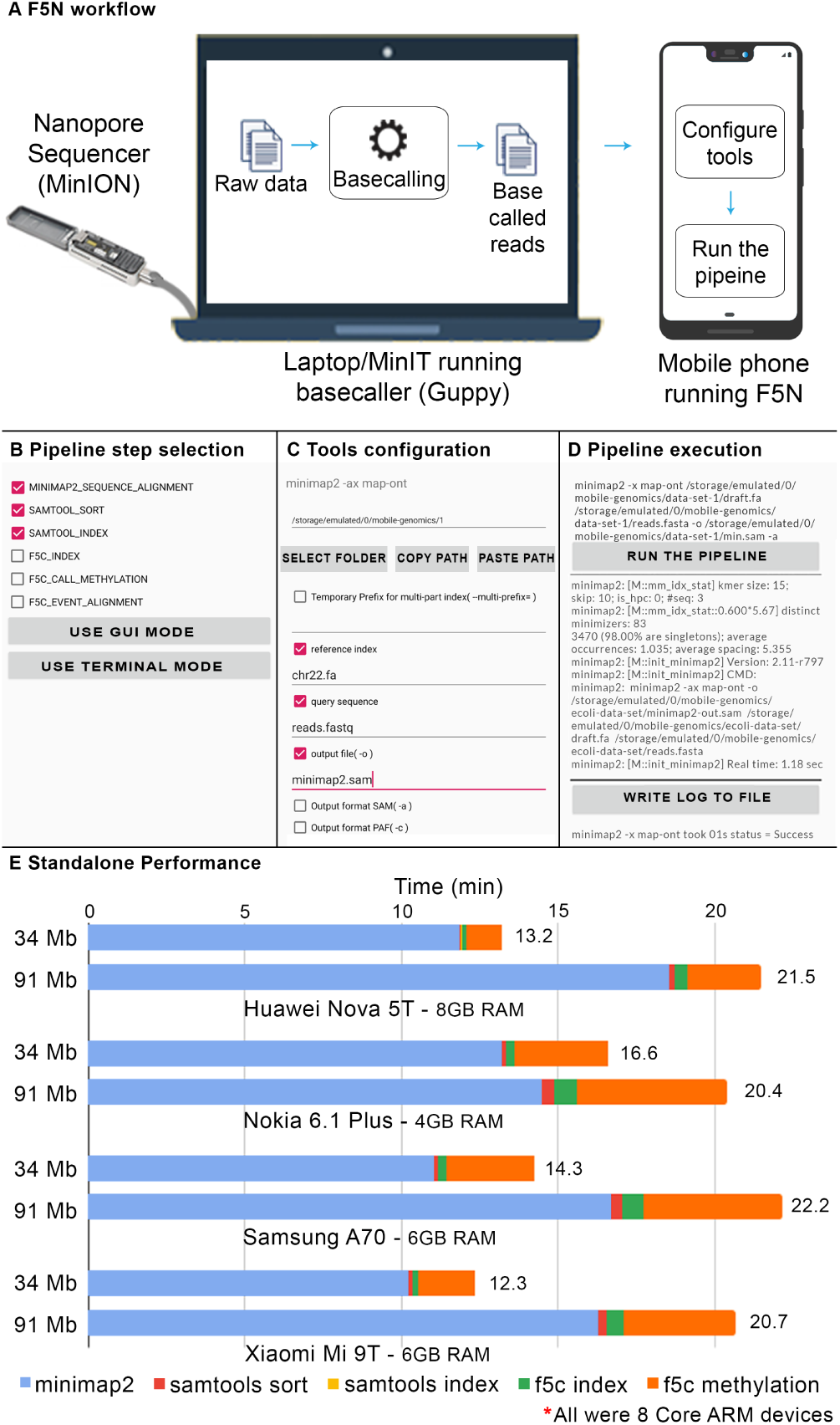
(A) Real-time analysis work setup. MinION connected basecaller and basecalled data transferred to a mobile device. (B,C,D) Screenshots of *F5N*. (E) Performance evaluation on four smartphones for two datasets (91 Mbases and 34 Mbases)

The ultra-portable MinION sequencer has proven to be beneficial for in-the-field sequencing in a variety of environments, such as Ebola surveillance in West Africa [2], microbial communities inspection in the Arctic [3] and DNA sequencing on the International Space Station [4]. The scarcity of portable and offline analysis solutions restricts the true potential of these sequencers. For instance, during Ebola surveillance, scientists encountered unexpected Internet connectivity issues where 3G signals dropped to 2G, massively increasing the data upload time. Mobile applications like *F5N* would be beneficial in such instances. In addition, mobile applications like *F5N* have the potential to increase the value of the ONT’s SmidgION, the upcoming smartphone pluggable sequencer.

*F5N* can execute an analysis process (known hitherto as the pipeline) on a nanopore dataset copied or downloaded by the user on to the phone storage. Out of the available tools *Minimap2, Samtools* and *F5C*, the user can select to run an individual tool, a combination of tools or the whole methylation detection pipeline (Fig. 1B). The intuitive graphical user interface in the application allows the user to configure the most common parameters for tools (Fig. 1C). A terminal environment is also provided in the application for an advanced user to provide command-line arguments. When the user starts the execution, the output is written to the phone storage as specified by the user and the log output is displayed on the application in real-time (Fig. 1D).

To get familiar with *F5N*, an example demonstration is provided inside the application which automatically downloads and executes a complete methylation calling pipeline on a small test dataset. *F5N* not only supports smaller genomes (eg: bacteria and virus) but also large genomes such as the human genome through an index partitioning approach as demonstrated in [5]. Refer Supplementary Sections II, III or the *Help* in the Android application on how to use *F5N*.

## Methodology

*F5N* Android Application (GUI and the framework) was developed using Java programming language. Popular long-read aligner *Minimap2* [6], the sequence data manipulator *Samtools* [7] and the methylation caller *F5C* [8] (optimised version of the popular tool *Nanopolish* [9]) were re-configured and cross-compiled to produce shared libraries (.*so* files) to run on Android over the ARM processor architecture. The interface between the Android Java application and the native code (compiled shared libraries) was written using the Java Native Interface (JNI). This interface invokes the native functions in the compiled libraries and captures the log output using *Unix Pipes*. The captured output is displayed on *F5N GUI*. Android Software Development Kit (SDK) and the Android Native Development Kit (NDK) were used as development tools.

One can re-configure different other bioinformatics tools written in C/C++ by following the detailed methodology in Supplementary Section VI and integrate them to *F5N* by following the guide in Supplementary Section VII. Thus, *f5c* is not just a toolkit, but also a framework for integrating existing or future C/C++ based command line bioinformatics tools. The challenges imposed by the restrictions in Android Operating system along with the methods to overcome them are discussed in Supplementary Section VIII.

## Benchmark Results

We benchmarked *F5N* using two publicly available NA12878 nanopore MinION datasets (flowcell IDs FAB42804 and FAF05869) [10]. A complete methylation calling pipeline was executed: read alignment to the full human reference genome (GRCh38) using *Minimap2* (8 index partitions), sorting and indexing of the alignments using *Samtools* followed by methylation calling using *F5C* (Supplementary Section I). Benchmarking was performed on four different phones (device specification in Supplementary Table S1). For full experiment details refer Supplementary Section III.

FAB42804 MinION dataset being comparatively smaller (only 16688 reads, 91.15 Mbases), *F5N* took 21.18 minutes in average to complete the whole methylation calling pipeline on it (Fig. 1E and Supplementary Table S2). A percentage of 77.8% of the execution time was consumed by *Minimap2* read alignment and 18.0% percentage of the time was consumed by *F5C* methylation calling. The highest and the second highest peak RAM were recorded for *F5C* methylation calling (3.4 GB) and *Minimap2* reads alignment (2.2 GB) respectively. By adjusting the memory governing parameters of each tool, peak RAM can be reduced to support a variety of low end devices (Supplementary Section IV).

FAF05869 MinION dataset (451,359 reads, 3.89 Gbases) was used to create batches of 4000 reads (∼34.49 Mbases per batch on average) to mimic the batch processing behaviour of the base-caller. Batches of reads were assigned on to all four mobile phones and were processed in parallel. A total time of 14.09 minutes was recorded in average for a single batch to complete the pipeline (Fig. 1E and Supplementary Table S4). A percentage of 89.8% of the execution time was for Minimap2 read alignment. The highest average peak RAM of 3.0 GB was recorded for *Minimap2* read alignment (Supplementary Table S5). Given a sequencing run on the MinION takes 48 hours, a single mobile phone would have been adequate to perform analysis on-the-fly for this particular dataset (with a power source to keep the phone charged) — one batch could be processed in 14.09 minutes while the sequencer would produce a batch only after every 25.52 minutes in average (see Supplementary Section V). Our future work will focus on harnessing the computing power of multiple mobile phones in parallel to keep-up with sequencing runs that produce larger data volumes.

## CONCLUSION

*F5N* demonstrates the true potential of portable genomics by executing a complete nanopore methylation calling pipeline locally on an Android mobile phone. A batch of 4000 nanopore reads (∼34 megabases) could be processed in ∼15 minutes where read alignment to the human genome, sorting and methylation calling consumed ∼12, *<*1 and ∼2 minutes, respectively. *F5N* can also be used by the community as a framework for integrating other tools and pipelines for nanopore data analysis. As future work, we will extend *F5N* to seamlessly connect multiple mobile phones to the base-calling device (laptop or ONT MinIT) through Wi-Fi, to process data even faster and on-the-fly during a sequencing run.

## Acknowledgments

Many thanks to Chandima Samarasinghe, Harshana Weligampola, Nirodha Suchinthana, James Ferguson, Rahal Medawatte, and Yasiru Ranasinghe for providing valuable feedback after testing *F5N*.

## Supplementary Materials

### I. Call methylation pipeline

**Fig. S1.**
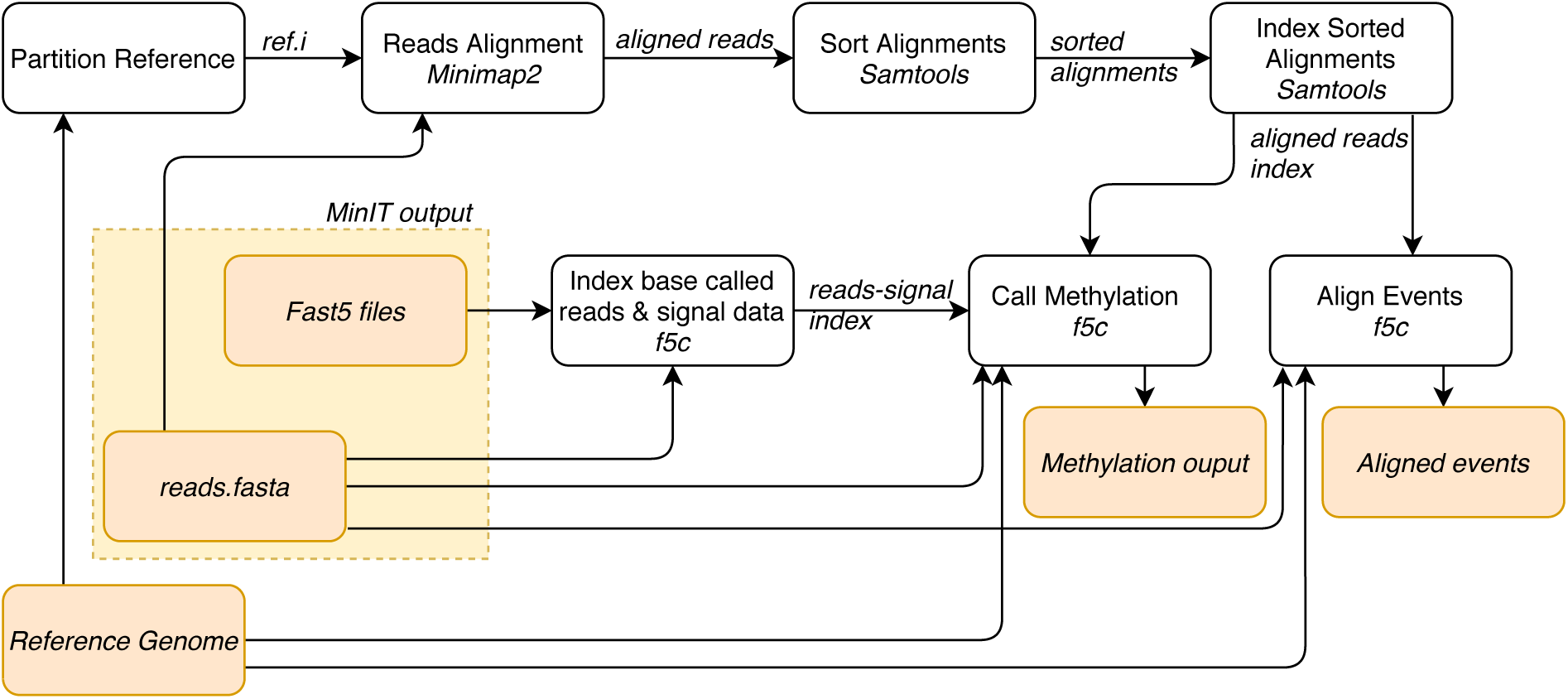
Methylation Calling (and Event Alignment) Pipelines comprise of three analysis tools - *Minimap2, Samtools* and *F5C*. Partition reference step is only required for larger genomes that do not fit in RAM.

*F5N* supports two complete DNA analysis pipelines - methylation calling and event alignment. Both pipelines are very similar except for the last steps. In this paper, we explain only the methylation calling pipeline. The pipeline has six steps (Fig. S1). The pipeline’s inputs are a reference genome, ONT raw signal data (.*fast5* files) and the corresponding base called reads (*reads.fastq(a)* file). The first step of the pipeline is to align base called reads to the reference genome. The widely used reads alignment tool is *Minimap2* [6] [11]. However, *Minimap2’s* memory usage grows with the size of the reference genome. Since a mobile phone has limited memory, *Minimap2* crashes for larger references. A novel algorithm was introduced to overcome the memory constraint using partitioned index [5]. We integrate this technique into our pipeline along with *Minimap2*. Please note that the reference partitioning should be done on a computer and the partitioned reference can be stored on a mobile phone (Supplementary Section III). Secondly, the aligned reads are sorted. Then these sorted results are indexed. Both sorting and indexing tasks use *Samtools* [7]. The final phase of the pipeline is polishing. *Nanopolish* [12] is the widely used polishing tool for nanopore data. We adopt a re-engineered version of *Nanopolish* called *F5C* [8], which is both memory and time efficient. *F5C* first indexes base called reads and raw signal data. Subsequently, *F5C* can either perform methylation calling or event alignment. In Event alignment pipeline, event alignment step is performed instead of methylation calling.

### II. F5N Guide

**Fig. S2.**
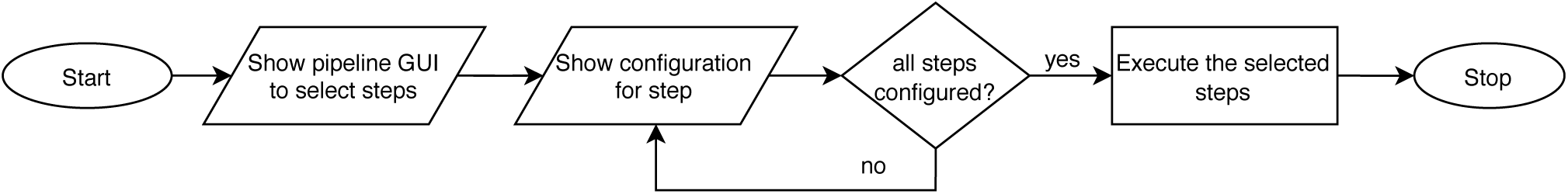
*F5N stand-alone mode* work flow

*F5N* has four major functionalities which are listed below.

1. *Stand-alone mode* for configuration and execution of a custom pipeline on the mobile device.
2. *Mobile-cluster mode* for real time analysis using a cluster of mobile devices, which is currently under development and will not be discussed in this paper.
3. Download data-sets using URLs and extract compressed files.
4. An example demonstration of downloading and extracting a nanopore data-set of E. Coli Bacteria, followed by executing a complete methylation calling pipeline on the data-set.

*F5N* includes a *help* section for new users to get started. It has a summary of the above four major functionalities (Fig. S3 B). Once *F5N* mobile application is launched for the first time, the user is prompted to grant permission to read and write from the internal storage of the mobile device. In most of the devices, once this permission is set, it is adequate to read from and write to the external storage (SD card) as well. However, in certain devices the user is expected to set this permission explicitly, which can be done by navigating to *[help → set SD card permission]* section (Fig. S3 C).

*F5N’s* start page has listed down the above four functionalities (Fig S3 A). A user navigating to *stand-alone mode* will land on a page to select custom pipeline steps from where he can choose *Minimap2, Samtools, F5C* or a desired combination of those tools (Fig. S3 F). Once the steps are selected, the user can choose either *GUI mode* or (Fig. S3 G and H) to configure parameters for each tool. Fig. S2 shows the procedure to use *stand-alone mode*. It is recommended to use the *GUI mode* as the final commands are always compiled into a set of strings and later shown in the *terminal mode* before proceeding to the execution. In *GUI mode* file path arguments get auto completed once the user set the correct path to the data set directory. *F5N* provides an elegant directory navigator for this purpose and both *GUI mode* and *terminal mode* have it. If the user chose *terminal mode* at the beginning, he skips *GUI mode* and lands on the *terminal mode*. From the *terminal mode* the user can proceed to the pipeline execution page (Fig. S3 I). Once the pipeline execution is started a timer will be displayed. After the execution of the pipeline the user can write results to a log file (named *f5n.log*) which is located inside *storage/mobile-genomics* directory. In the rare event of a crash of *F5N*, the user can run the previous pipeline using *LOAD PREVIOUS CONFIGURATION* command. If the app crashes during an execution, the user can identify the error occurred by referring *tmp.log* which is located inside *storage/mobile-genomics* folder. For more information regarding log files please refer the *help* section on *home page* in the application.

Functionalities to download a data-set form a URL and extract a compressed data-set are available on a same page (Fig. S3 D). To download a data-set, the user has to set the specific data-set URL path and the location on the storage to where the data-set should be downloaded. Decompressing a file is as easy as setting the file path of the compressed file and pressing the *EXTRACT* button. The decompressed file will have the same location as the compressed file’s. Since a data-set usually consists of many numbers of considerably small .*fast5* files, it will take much time to transfer them to a device storage unless the files are compressed. Hence, *F5N* is provided with a file extraction functionality to decompress the files as necessary. In *mobile-cluster mode* compressed files will get transferred over WiFi.

The example demonstration is a setup with only three steps to help users get familiar with *F5N*. The steps involve the basic procedure to execute a pipeline. They are 1, download a data set 2, extract the data set and 3, execute the pipeline (Fig. S3 E).

**Fig. S3.**
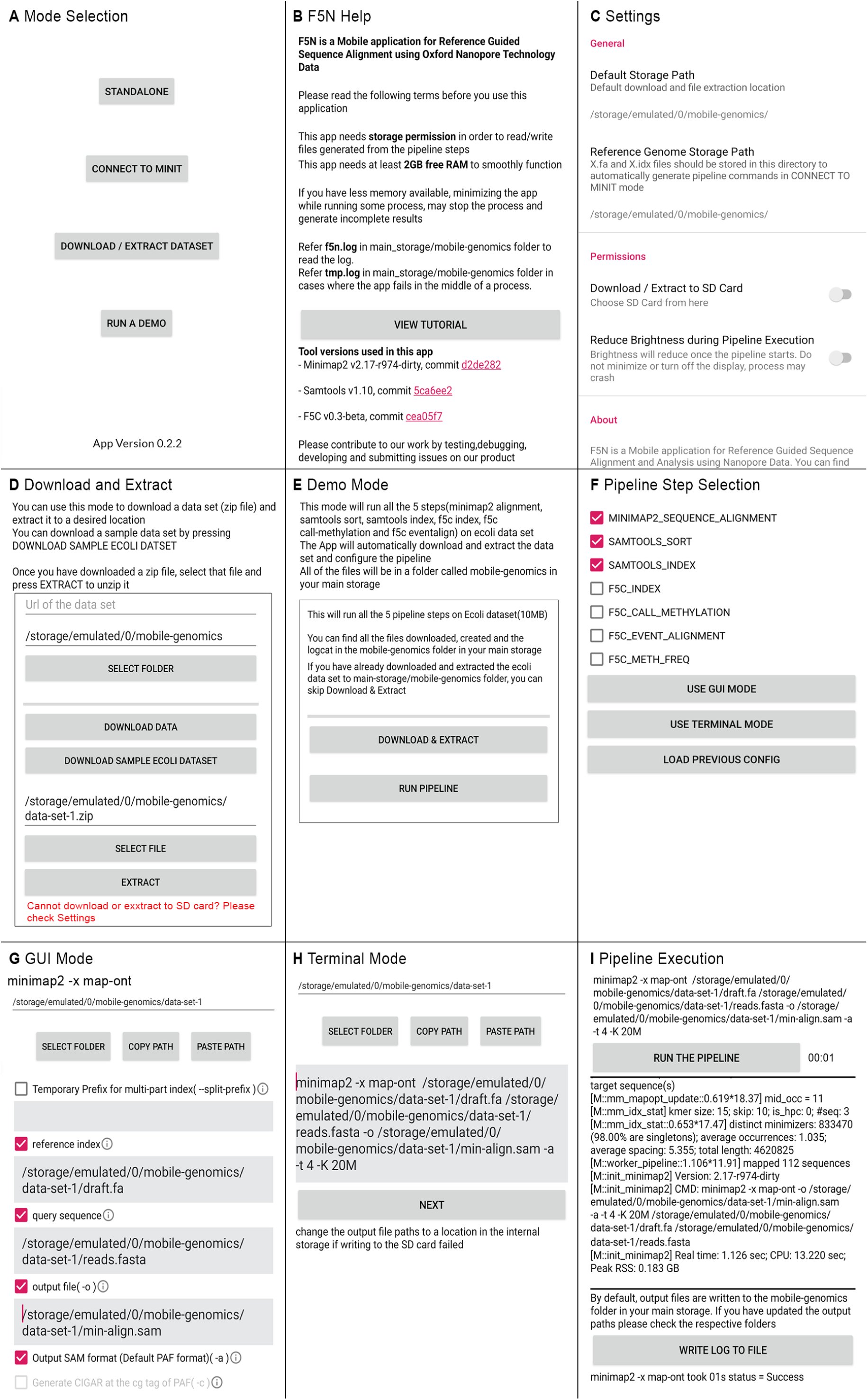
*F5N* screenshots (A) Homepage. (B) Help page. (C) Set F5N Settings. (D) Download and extract files. (E) Run an example pipeline. (F) Select pipeline steps. (G) Configure pipeline steps using GUI. (H) Configure pipeline steps using terminal environment. (I) Pipeline execution

### III. Detailed F5N examples and results

In this section, we present a detailed overview of the tests run using *F5N*. GRCh38 human genome reference was used as the reference (3.1 GB). For reads alignment, the index constructed for the whole reference by *Minimap2* is too large to fit in RAM (∼8 GB). It was partitioned into 8 partitions using the script [13]. The resulting partitioned index is ∼8.2 GB in size. Since now only a single partition is loaded to RAM at once, the memory of a mobile device is sufficient for *Minimap2* alignment. User has to store both the reference genome and its partitioned version either on the internal storage or on the external storage (SD card). Please refer Supplementary Section VIII on how to choose the SD card format. Two datasets, flowcell id-FAB42804 and flowcell id-FAF05869 were downloaded from the publicly available NA12878 nanopore sequenced data repository [10].

The first dataset (FAB42804) had 16688 reads (91.15 Mbases) and the second dataset had 451,359 reads (3.89 Gbases). The first dataset was used as is to execute a complete methylation calling pipeline. To mimic the batch processing behaviour of the basecaller, the second dataset (FAF05869) was divided into batches of 4000 reads (∼34.49 Mbases per batch on average). For this, the original raw signal files (single-fast5 files) of FAF05869 were downloaded and converted to multi-fast5 using ONT’s *single_to_multi_fast5* tool (version 1.4.8) with default options. Then, these multi-fast5 files were re-basecalled using ONT’s basecaller *guppy* (version 3.3.0) under the *dna_r9.4.1_450bps_hac* model.

The devices listed in the table S1 were used in the experiments. All the devices were octa-core arm64-v8a instruction set devices. Methylation calling pipeline illustrated in the Supplementary Section I was executed on the first dataset and on batches from the second dataset (2 batches per device). The datasets along with the reference and partitioned index were copied to mobile devices through a USB connection. Each test was repeated 3 times on a device and the tables S2-S5 show the average values. The results are discussed in the paper.

Below, are the commands ran on a device to execute the complete methylation pipeline. These commands were generated through *F5N’s GUI mode*, except for the first command.

1. *Partitioning the reference genome (on a computer)*

~~~
divide_and_index.sh [reference .fa file] [num_parts [8]] [output .idx file] [min-
imap2_exe] [minimap2_profile [map-ont]]
~~~
2. *Aligning reads to the partitioned human genome using Minimap2*

~~~
minimap2 -x map-ont --split-prefix [temporary file path] [partitioned reference]
[reads.fastq|fasta] -o [output .sam file] -a -t [no. of threads [4]] -K [Number of bases
loaded into memory to process in a mini-batch [5M]]
~~~
3. *Sort aligned reads using Samtools*

~~~
samtool sort [minimap2 output] -o [output .bam file]
~~~
4. *Index sorted reads using Samtools*

~~~
samtool index [samtools sort output]
~~~
5. *Index fast5 files*

~~~
f5c index --directory [fast5_folder] [reads.fastq|fasta]
~~~
6. *DNA methylation calling using F5C*

~~~
f5c call-methylation -r [reads.fastq|fasta] -b [path to samtools index output] -g [ref.fa]
-o [output meth.tsv] -B [max number of bases loaded at once [2.0M]] -K [max number of reads
loaded at once [256]]
~~~
7. *Aligning nanopore events to reference k-mers using F5C (optional, not a step in methylation calling)*

~~~
f5c eventalign -r [reads.fastq|fasta] -b [path to samtools index output] -g [ref.fa]-o
[output meth.tsv] -B [max number of bases loaded at once [2.0M]] -K [max number of reads
loaded at once [256]] --summary [output events.summary.txt]
~~~

**TABLE S1.** List of mobiles used in the experiments

| Device | Manufacturer | Model | RAM(GB) | Internal Storage(GB) |
| --- | --- | --- | --- | --- |
| A | Huawei | Nova 5T | 8 | 128 |
| B | Nokia | 6.1 Plus | 4 | 64 |
| C | Samsung | A70 | 8 | 120 |
| D | Xiaomi | Mi 9T | 6 | 120 |

**TABLE S2.** F5N performance for the complete NA12878-FAB42804 dataset (91 Mbases)

| Device | Minimap2 (min) | Samtools sort (min) | Samtools index (min) | F5C index (min) | F5C call methylation (min) | Total time (min) |
| --- | --- | --- | --- | --- | --- | --- |
| A | 18.52 | 0.20 | 0.02 | 0.37 | 2.37 | 21.48 |
| B | 14.48 | 0.38 | 0.02 | 0.70 | 4.82 | 20.40 |
| C | 16.67 | 0.36 | 0.03 | 0.65 | 4.45 | 22.68 |
| D | 16.27 | 0.28 | 0.02 | 0.53 | 3.58 | 20.18 |

**TABLE S3.** Peak RAM of individual steps when processing NA12878-FAB42804 dataset

| Device | Minimap2 (GB) | Samtools sort (GB) | Samtools index (GB) | F5C index (GB) | F5C call methylation (GB) |
| --- | --- | --- | --- | --- | --- |
| A | 2.20 | 0.32 | 0.19 | 0.15 | 3.95 |
| B | 2.22 | 0.26 | 0.09 | 0.10 | 2.78 |
| C | 2.27 | 0.30 | 0.14 | 0.15 | 3.58 |
| D | 2.22 | 0.34 | 0.18 | 0.19 | 3.28 |

### IV. Adjusting memory governing parameters of the tools

The following steps can reduce peak RAM usage, to run *F5N* on a broad spectrum of mobile devices. For more details visit *Minimap2* man page[14], *Samtools* man page[15] and *F5C* man page[16].

**TABLE S4.**
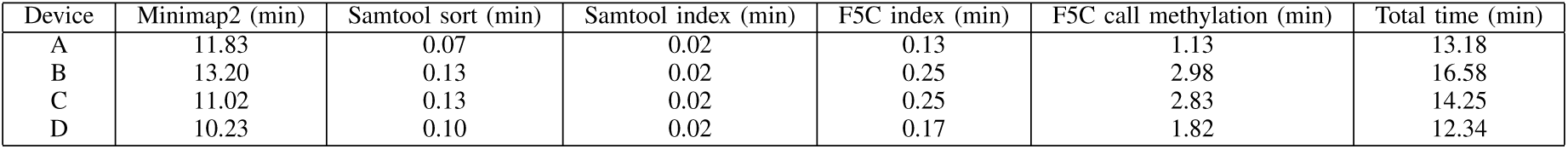
F5N performance for a batch of 4K reads (34 Mbases) from NA12878-FAF05869 dataset

| Device | Minimap2 (min) | Samtool sort (min) | Samtool index (min) | F5C index (min) | F5C call methylation (min) | Total time (min) |
| --- | --- | --- | --- | --- | --- | --- |
| A | 11.83 | 0.07 | 0.02 | 0.13 | 1.13 | 13.18 |
| B | 13.20 | 0.13 | 0.02 | 0.25 | 2.98 | 16.58 |
| C | 11.02 | 0.13 | 0.02 | 0.25 | 2.83 | 14.25 |
| D | 10.23 | 0.10 | 0.02 | 0.17 | 1.82 | 12.34 |

**TABLE S5.** Peak RAM of individual steps when processing a batch of 4K reads from NA12878-FAF05869 dataset

| Device | Minimap2 (GB) | Samtool sort (GB) | Samtool index (GB) | F5C index (GB) | F5C call methylation (GB) |
| --- | --- | --- | --- | --- | --- |
| A | 2.48 | 0.16 | 0.09 | 0.10 | 0.99 |
| B | 3.16 | 0.22 | 0.14 | 0.16 | 1.18 |
| C | 3.16 | 0.22 | 0.14 | 0.16 | 1.18 |
| D | 3.41 | 0.23 | 0.16 | 0.17 | 1.19 |

1. Increase the number of partitions the genome reference is split into, to reduce peak RAM usage in *Minimap2 alignment*
2. In *Minimap2 alignment* reduce the parameter value, *number of bases loaded into memory to process in a mini-batch [-K]*
3. In *Samtools sort* reduce the parameter value, *the maximum required memory per thread [-m]*
4. In *F5C call methylation* reduce the parameter value, *batch size (max number of reads loaded at once) [-K]*
5. In *F5C call methylation* reduce the parameter value, *max number of bases loaded at once [-B]*
6. In *F5C call methylation* skip ultra long reads by setting the option *[–skip-ultra FILE]*
7. In *F5C call methylation* reduce the threshold value to skip ultra long reads *[–ultra-thresh INT]*

### V. Caclulating base-calling rate

Given the total number of reads in a dataset *N*, the number of reads in a base-called batch *B* and the duration of sequencing run *T*, we can calculate the average time taken by the basecaller to produce a batch (base calling rate) *R* using the following equation, 

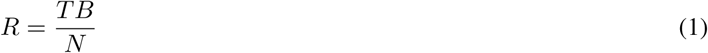

In our case *T* = 48 hours, *B* = 4000 and *N* = 451,359 which results in *R* to be 25.52 min.

### VI. Reconstructing sequence analysis tools for ARM/Android

**Fig. S4.**
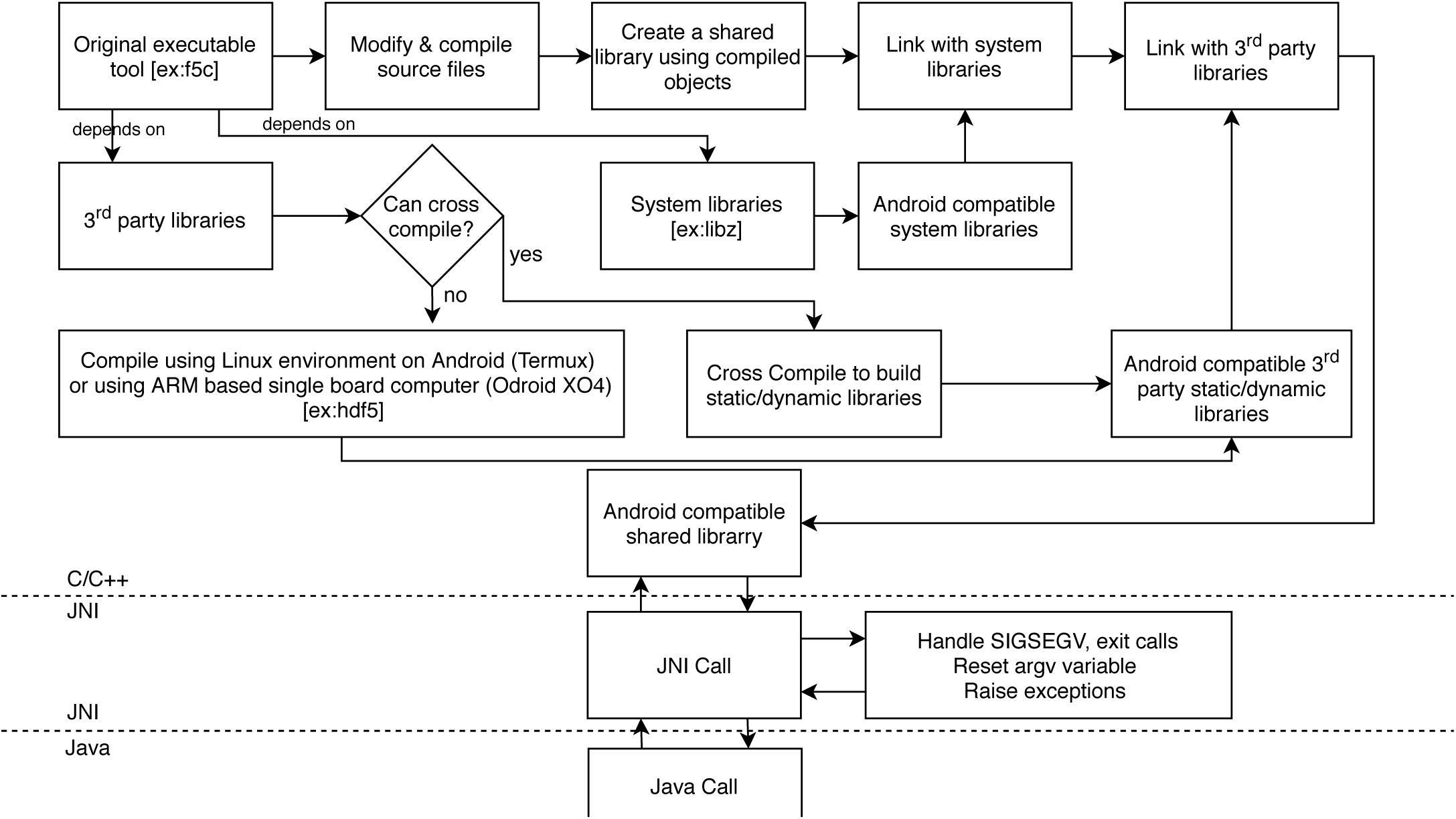
Cross-compiling work flow and JNI interface design involved in including a new sequence analyis tool to *F5N*.

#### A. Introduction and prerequisites

This section provides fundamental guidelines on how to compile an analysis tool written in C/C++ for Android. Except for a few essential amendments (discussed below), our approach does not alter the original C/C++ code, which permits straightforward integration of new C/C++ analysis tools into *F5N*.

Having some prior experience with Android SDK (Software Development Kit), Android Studio[17], NDK (Native Development Kit) [18], Java Native Interface (JNI)[19], ADB (Android Debug Bridge)[20], CMake build manager[21] and Ninja build system[22] certainly helps someone who is interested in rebuilding *F5N* or extending it with new tools.

#### B. Native compiling an analysis tool

In our case, the fact that all the necessary tools (*Minimap2, Samtools* and *F5C*) were written in C/C++ let us to follow the same approach to compile them for Android. The approach can be summarized as shown in Fig. S4. Before starting the compilation we tested the tools using *Termux* (a terminal emulator with Linux environment for Android)[23]. In *Termux*, the tools’ git repositories were cloned and built as instructed on the respective installation guides. This method can be known as *native compiling*. If a tool gets *natively compiled* in *Termux*, we can reasonably expect the tool to get cross-compiled for Android. There are two major *Instruction Set Architectures (ISA)* for *ARM* mobile devices called *armeabi-v7a and arm64-v8a. Termux* was installed on multiple devices with different architectures to make sure that the tools can be *natively compiled* in both the architectures. It is noteworthy that in *Termux*, the executable versions (.*exe* format) of the tools were built in contrast to their dynamic library versions (.*so* format). To build an Android application, the dynamic version of an original tool is required[24]. All the Android applications are a subset of Java programs and the gateway to the native (C/C++) code is obtained by loading the dynamic version of the native tool. To obtain a dynamic version of the tool (.*so* format), we can change the build configuration and natively compile in *Termux*. However, this method is not encouraged as it can impose device based restrictions on .*so* files which can result in intensive and costly debugging. The recommended method to obtain the dynamic version of a tool is to cross-compile using the Android Tool-chain[25]. This eliminates the burden associated with *native compiling* and simplifies the process of updating the tools to their latest versions. This in return automates continuous integration and delivery. In Suppliementary Sections VI-E and VII we provide further details on how to cross-compile.

#### C. Determining third party libraries used by an analysis tool

It is necessary to determine the third party libraries used by the analysis tools (*Minimap2, Samtools* and *F5C*). A tool to function correctly, associated third party libraries should also be linked statically or dynamically with the tool, i.e., third party libraries should also be cross-compiled. *Samtools* depends on *htslib* [26] library. F5C depends on both *htslib* and *HDF5* [27] libraries. Hdf5 library is used to handle fast5 files and there exists no straightforward method to cross compile hdf5. Hence we used native compiled instances of *HDF5* library (the instances that were built using Termux). We maintained two instances of *HDF5*, one for each Architecture (*armeabi-v7a* and *arm64-v8a*). These third party libraries were statically linked to respective shared libraries. If a third party library is being used by two dependent tools, make sure to link a dynamic version of the library. For an example in our case, *htslib* was used by *Samtools* and *F5C*. Trying to statically link *htslib* separately to each tool, caused the software to crash on some devices. This was resolved by linking *htslib* dynamically.

#### D. Designing JNI interface

JNI acts as the bridge between native (C/C++) methods and Java function calls. In JNI interface, each tool’s *int main(int argc, char* argv[])* function was called. The function name, *int main(int argc, char* argv[])* was renamed as *int init_X(int argc, char* argv[])* where X was *Minimap2,Samtools* or *F5C*. This function renaming is necessary as the tools are not stand-alone executable applications but dynamic libraries. Moreover, a header file was introduced for each tool that contained the function signature *int init_X(int argc, char* argv[])*. JNI interface was extended to facilitate the following,

1. Handle *exit* signals returned by native code.
2. Handle *SIGSEGV* signals returned by native code.
3. Raise exceptions on behalf of the native functions.
4. Reset *argv* variable before calling another native function [*int init_X(int argc, char* argv[])*].
5. A tunnel between the native code and the Java program to communicate *standard error* messages.
6. If original code does not define an output file path argument, redirect *standard ouput* to a file.

It is important to handle different types of signals and errors thrown by the native code to prevent JVM from crashing. For example the original code most probably will exit with an error if something goes wrong. This kills the JVM if not handled. We want to keep the JVM running throughout a *F5N* session. In order to do that, the exit call should be caught and handled in JNI [28]. In a similar manner, *SIGSEGV* signals should be handled safely [29]. Once an exit call or *SIGSEGV* signal is handled this should be informed to Java program so that the user can investigate the problem. To do this, exceptions are thrown from JNI to Java [30].

The original C/C++ tools take input as command line arguments. When a pipeline with more than one step is executed, the native code attempts to read the same argument vector multiple times. If the argument vector is not reset after the completion of the first step, arguments do not get parsed in the second step as desired. This resetting part is not implemented in most of the original libraries. In JNI this should be implemented to avoid failures associated with arguments parsing [31].

**Fig. S5.**
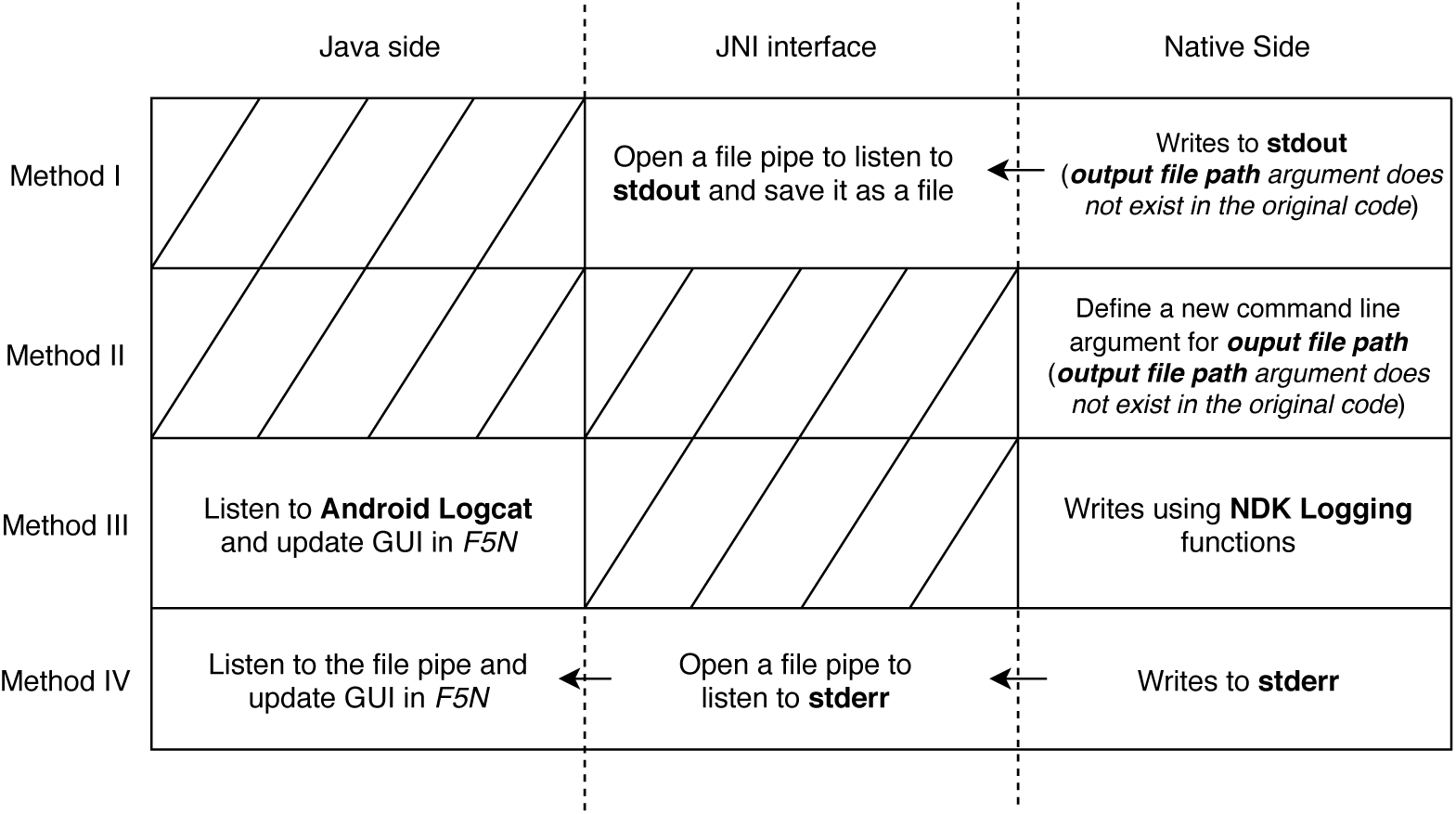
Method I - Using a file pipe to listen and save *standard output*. Method II - Define a new command line argument for *output file path*. Method III - Using NDK Logging functions to print *standard error* to Logcat. Method IV - Using a file pipe to listen *standard error*.

Typically a tool writes the results to the *standard ouput* and meta-information to the *standard error*. On Android, we want the results to be written to a file and meta-information to be displayed on a GUI in real-time. To write results to a file, most of the tools provide a command line argument called *file output path*. Otherwise, in terminal environments like in Linux, we can redirect the *standard output* to a file (using the output redirection operator ‘*>*‘). On Android, this is not possible. To overcome this issue two methods can be adopted. The first is to open up a file pipe in JNI to listen to the *standard ouput*, where the results will get written to this file (Fig. S5 Method I). The second is to define the output file path as an argument in the original code (Fig. S5 Method II). However, all the libraries in *F5N* had output file path as an argument. Now we present two methods to catch the *standard error*, which should be displayed to the user in real-time. The first method is to replace all the *fprintf(stderr,…)* functions with functions defined in NDK logging [32]. Then the *standard error* will get written to Android Logcat [33]. From Android Logcat, it is again piped to be displayed (Fig. S5 Method III). In practice, this method does not guarantee to display the complete set of messages written to the *standard error* during an execution. The more robust method is to open a file descriptor to listen to the *standard error*. Then this file should be read in Java side and the GUI should be updated (Fig. S5 Method IV). This involves no amendments in the original libraries but a declaration of file descriptors in *JNI* code.

#### E. Cross-compiling an analysis tool

To facilitate the modifications discussed above, the original repositories of the tools were forked and changed. Different tools use different build configurations. The build scripts for each tool were re-written using CMake. Suppose an original tool is built with *GNU Make* using a *Makefile*. In that case, one has to extract the *source files, header files, compiler flags, linked libraries etc* to create a *CMakeLists.txt* file. Refer how a *CMakeLists.txt* was written for *htslib* [34]. CMake along with Ninja is the recommended native build setup for Android. Compiling with CMake allows ADB to go deep into the native code when debugging. This was really helpful to figure out the static/dynamic version issue related to *htslib* library. It was straightforward to link the necessary third party libraries(*HDF5* and *htslib*) and system libraries (*libz, liblog, libm* etc) with original libraries using CMake. To follow the full set of modifications please refer *Minimap2* [35] *Samtools* [36] and *F5C* [37]. One can build libraries without using CMake but by using already available *Standalone Toolchains*[38]. However, this eliminates the possibility to debug using ADB.

#### F. Dynamic object construction in Java side

The part of *F5N* written in Java is dynamic and adaptive. For example the arguments set for each tool is stored in JSON format and objects are created by converting JSON objects to Java objects. In this way, *F5N* and arguments are decoupled making it easy to alter the format of the arguments if needed. The widgets linked with arguments are drawn programmatically instead of manually drawing them on the layout. This makes it easy to extend *F5N* with new analysis tools (refer Supplementary Section VII).

### VII. Integrating a new analysis tool to F5N

The following steps summarize the work flow to add a new DNA analysis tool written in C/C++ to *F5N*. Please refer Fig. S6 for *F5N* directory structure. *F5N* repository is available at https://github.com/SanojPunchihewa/f5n

**Fig. S6.**
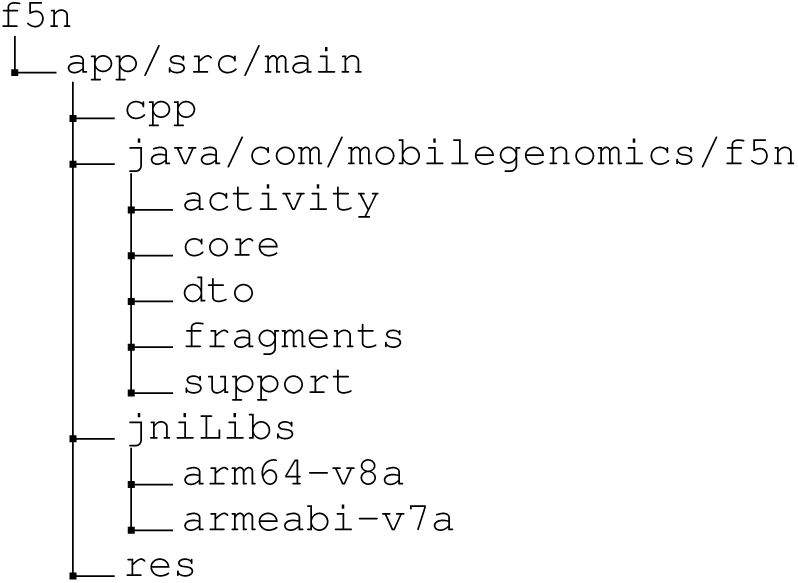
*F5N* project directory structure, only the important folders are shown. The directory *cpp* contains *CMakeLists.txt* and *interface_X.h* header files. The .*so* files are stored according their *ISA* inside *jniLibs* directory. The Java Class *PipelineStep* is inside directory *core*.

1. Identify third party libraries that the new tool depends on. e.g. *Samtools* depends on *htslib*.
2. Create dynamic versions of third party libraries if they are used by other analysis tools in *F5N*.
3. Create static versions of third party libraries if they are not used by other analysis tools in *F5N*.
4. Cross compile the new tool and link it with static third party libraries to create a dynamic library.
5. Place all the dynamic libraries in *jniLibs/[ANDROID_ABI]* directory.
6. Repeat the above steps for different *ANDROID_ABIs*. (*armeabi-v7a and arm64-v8a*)
7. Change *CMakeLists.txt* file in *cpp* directory to include dynamic third party dependencies.
  - ~~~
add_library(libnewdependency SHARED IMPORTED)
~~~
  - ~~~
set_target_properties(libnewdependency PROPERTIES IMPORTED_LOCATION
CMAKE_SOURCE_DIR/../jniLibs/$ANDROID_ABI/libnewdependency.so)
~~~
8. Change *CMakeLists.txt* to include the new analysis library.
  - ~~~
add_library(libnew SHARED IMPORTED)
~~~
  - ~~~
set_target_properties(libnew PROPERTIES IMPORTED_LOCATION
CMAKE_SOURCE_DIR/../jniLibs/$ANDROID_ABI/libnew.so)
~~~
9. Change *CMakeLists.txt* file to link the new analysis library.
  - ~~~
target_link_libraries(native-lib libminimap libsamtool libf5c
libhts libnew libnewdependency1 libnewdependency2 … $log-lib)
~~~
10. Copy *interface_X.h* file with the function signature *init_X(int argc, char* argv[])* to *cpp* directory (X is the name of the new library).
11. Create *enum* entries in *PipelineStep* Class located in *core* directory for the sub-tools (commands) in the new analysis tool.
  - ~~~
NEW_TOOL_NAME(COMMAND_ID, “tool_name sub-tool_name(command)”);
ex:_F5C INDEX(3, “f5c index”);
~~~
12. For each sub-tool (command) create a *JSON* file containing the list of arguments and their default values[39].
13. Link JSON files to the *configureArguments* method in *GUIConfiguration* Class located inside *java/com/mobilegenomics/f5n* directory, i.e. add new switch cases to match *NEW_SUB_TOOLs* and assign *JSON* files appropriately.
14. Import *interface_X.h* header file to *native-lib.cpp* source file located inside *cpp* directory.
  - ~~~
include “#interface_X.h”
~~~
15. Follow a similar approach to other tools to call *init_X(int argc, char* argv[])* function in *native-lib.cpp* source file.

Moreover, the following list comprises of common mistakes that could happen when extending or rebuilding *F5N*. Please refer Supplementary Section VI for details.

1. Overlooking compiler flags when creating a CMake build configuration. e.g. not including the compiler flag *-D_FILE_OFFSET_BITS=64* in *Minimap2* CMake build will result in file read failures.
2. Statically linking third party libraries that are used by more than one tool. e.g. Linking the static version of *htslib* library caused *Samtools* and *F5C* to fail.
3. Not Handling native exceptions in JNI interface
4. Not Handling native SIGSEGV signals and exit signals in JNI interface
5. Not Resetting command line argument vector before calling a new native function

### VIII. Advanced details

*F5N* is not built as an Android background process. That is because when processing a dataset, *F5N* may consume memory more than the recommended amount for a background process. Android kills such over memory consuming background processes. Hence, *F5N* is built as a regular Android application. However, this introduced a caveat. That is when running a pipeline, the device display should be kept on. That is because Android interrupts running applications once the display goes off. Hence, *F5N* has to keep the display on and this increases power consumption. Our workaround is to reduce the display brightness to its minimal value, once a pipeline execution starts. With this method, device B’s battery (3060 mAh) drains approximately by 214.2 mAh for a complete methylation calling pipeline on a batch data-set (∼34.49 Mbases on average).

Android tends to kill a process or destroy an activity if something goes wrong. In *F5N* usually it the case when the native code tries to over consume memory. Since this kill signal comes from the Android Kernel, it cannot be handled. The simple solution is to be aware of the device memory and set memory governing parameters accordingly (see Supplementary Section IV). Saving the state of the application, i.e, saving previously executed command and loading it later, saves time for re-configuring the pipeline (Fig. S3F). In the next attempt, the user is advised to tweak memory parameters as it is the solution most of the time.

Already executing native code on Android cannot be stopped arbitrarily. The ramification of this is that the user cannot suspend or stop an executing pipeline. On the other hand, JVM being a multi-threaded process and JNI calls not being *POSIX async-signal-safe*, it is not possible to run the native code in a cloned process. As a consequence peak RAM usage will record the highest RAM usage for the whole application session rather than for the latest pipeline execution. To circumvent both problems the “safest” solution is to restart *F5N*. An advanced method to suspend or stop an executing command is by adding interrupt listeners to the original code. That is while the original code is being executed, it periodically checks its environment for an interrupt signal, e.g, state of a flag value in a file. The flag value can be changed on-the-fly to stop or suspend the execution. Please note that in *F5N*, when a pipeline is running the user can still go to the previous activity. In such attempts, a warning message is displayed saying that the pipeline will stop. This does not guarantee the complete termination of the native process but the termination of the running Java thread. Hence, the user is advised to restart the application to safely terminate a pipeline.

A mobile phone has two storage types - internal storage and external storage (SD card storage). *FAT32* is a popular file system format used in SD cards. However, *FAT32* does not support files with more than the size of ∼4GB. Usually, the partitioned genome reference file exceeds the size 4GB. Therefore, the user is advised to use file system formats like *ext3, ext4, exFAT32* etc as the SD card file system format. Out of these formats, *exFAT32* is recommended. It is noteworthy that Google has introduced (from Android 8.0) a virtual file system wrapper called *SDCardFS* to regulate SD card access by Android applications. However, still, the SD card should have one of the compatible file system formats to work with larger files.

Downloading and extracting a dataset to the SD card can be done after setting SD card permission (refer Fig S3C). On Android, writing to the SD card storage via native code (C/C++) can only be done using Storage Access Framework (SAF)[40]. To implement SAF, most of the original code has to be changed. One workaround is to use the *Method I* as illustrated in Fig. S5. In JNI interface, we can use SAF to write the *standard output* to the SD card. However, there are sub-tools that directly write to files, e.g., *F5C index* writes files automatically to the dataset directory. In such scenarios, the original tool should be reconfigured to facilitate SAF. The current version of *F5N* does not support this feature. Therefore the user cannot perform writes to the SD card but can read from it.

## References

[1] Hengyun Lu, Francesca Giordano, and Zemin Ning. “Oxford Nanopore MinION sequencing and genome assembly”. In: Genomics, proteomics & bioinformatics 14.5 (2016), pp. 265–279.

[2] Joshua Quick et al. “Real-time, portable genome sequencing for Ebola surveillance”. In: Nature 530.7589 (2016), pp. 228–232.

[3] Jacqueline Goordial et al. “In situ field sequencing and life detection in remote (79 26’ N) Canadian high arctic permafrost ice wedge microbial communities”. In: Frontiers in microbiology 8 (2017), p. 2594.

[4] Sarah L Castro-Wallace et al. “Nanopore DNA sequencing and genome assembly on the International Space Station”. In: Scientific reports 7.1 (2017), pp. 1–12.

[5] Hasindu Gamaarachchi, Sri Parameswaran, and Martin A Smith. “Featherweight long read alignment using partitioned reference indexes”. In: Scientific reports 9.1 (2019), p. 4318.

[6] Heng Li. “Minimap2: pairwise alignment for nucleotide sequences”. In: Bioinformatics 34.18 (2018), pp. 3094–3100.

[7] Heng Li et al. “The Sequence Alignment/Map format and SAMtools”. In: Bioinformatics 25.16 (June 2009), pp. 2078–2079.

[8] Hasindu Gamaarachchi et al. “GPU Accelerated Adaptive Banded Event Alignment for Rapid Comparative Nanopore Signal Analysis”. In: bioRxiv (2019), p. 756122.

[9] Jared T Simpson et al. “Detecting DNA cytosine methylation using nanopore sequencing”. In: Nature methods 14.4 (2017), p. 407.

[10] Miten Jain et al. “Nanopore sequencing and assembly of a human genome with ultra-long reads”. In: Nature biotechnology 36.4 (2018), p. 338.

## Supplementary References

[11] Heng Li. “Minimap and miniasm: fast mapping and de novo assembly for noisy long sequences”. In: Bioinformatics 32.14 (2016), pp. 2103–2110.

[12] Jared Simpson. Nanopolish. https://github.com/jts/nanopolish.

[13] Hasindu Gamaarachchi. Index construction with chromosome size balancing. https://github.com/hasindu2008/minimap2-arm/tree/master/misc/idxtools.

[14] Heng Li. Manual Reference Pages - Minimap2 (1). https://lh3.github.io/minimap2/minimap2.html.

[15] Samtools. Manual page from samtools-1.10. http://www.htslib.org/doc/samtools.html.

[16] Hasindu Gamaarachchi. Manual Reference Pages - F5C. https://hasindu2008.github.io/f5c/docs/overview.

[17] Android. Android Studio and SDK. https://developer.android.com/studio.

[18] Android. Android NDK. https://developer.android.com/ndk.

[19] Android. JNI. https://developer.android.com/training/articles/perf-jni.

[20] Android. Android Debug Bridge. https://developer.android.com/studio/command-line/adb.

[21] Kitware. CMake build manager. https://cmake.org/.

[22] Ninja. Ninja build system. https://ninja-build.org/.

[23] Fredrik Fornwall. Termux Linux environment emulator. https://play.google.com/store/apps/details?id=com.termux&hl=en.

[24] Ian F Darwin. Android Cookbook: Problems and Solutions for Android Developers. “O’Reilly Media, Inc.”, 2017, “661–666”.

[25] Android. Android Cmake cross-compilation. https://developer.android.com/ndk/guides/cmake.

[26] samtools. htslib. https://github.com/samtools/htslib.

[27] Mike Folk, Albert Cheng, and Kim Yates. “HDF5: A file format and I/O library for high performance computing applications”. In: Proceedings of supercomputing. Vol. 99. 1999, pp. 5–33.

[28] JNI handle exit calls. http://jnicookbook.owsiak.org/recipe-no-016/.

[29] JNI handle SIGSEGV calls. http://jnicookbook.owsiak.org/recipe-no-015/.

[30] JNI throw exceptions. http://jnicookbook.owsiak.org/recipe-no-019/.

[31] Sanoj Punchihewa. F5N JNI interface. https://github.com/SanojPunchihewa/f5n/blob/master/app/src/main/cpp/native-lib.cpp.

[32] Android. Logging Android NDK. https://developer.android.com/ndk/reference/group/logging.

[33] Android. Logcat command-line tool. https://developer.android.com/studio/command-line/logcat.

[34] HTSLIB. CMake build for HTSLIB. https://github.com/hiruna72/htslib/blob/76f9eaa29a23573a70e37ca6ed842719e03cde55/INSTALL#L101.

[35] Minimap2. CMake build for Minimap2. https://github.com/SanojPunchihewa/minimap2-arm/tree/build-cmake#cmake-build.

[36] Samtools. CMake build for Samtools. https://github.com/hiruna72/samtools/blob/947c5b66cf91abc9b3e58b61642994dd8f4ae7e4/INSTALL#L103.

[37] F5C. CMake build for F5C. https://github.com/hiruna72/f5c/tree/cmake_build#building.

[38] Android. Standalone Toolchains (Obsolete). https://developer.android.com/ndk/guides/standalone_toolchain.

[39] F5N. F5N arguments JSON format. https://github.com/SanojPunchihewa/f5n/blob/master/app/src/main/res/raw/minimap2.json.

[40] Android. Open files using storage access framework. https://developer.android.com/guide/topics/providers/document-provider.html.

